# Developing a transcriptomic framework for testing testosterone-mediated handicap hypotheses

**DOI:** 10.1101/814178

**Authors:** Daniel J. Newhouse, Ben J. Vernasco

## Abstract

Sexually selected traits are hypothesized to be honest signals of individual quality due to the costs associated with their maintenance, development, and/or production. Testosterone, a sex steroid associated with the development and/or production of sexually selected traits, has been proposed to enforce the honesty of sexually selected traits via its immunosuppressive effects (i.e., the Immunocompetence Handicap Hypothesis) and/or by influencing an individual’s exposure/susceptibility to oxidative stress (i.e., the Oxidation Handicap Hypothesis). Previous work testing these hypotheses has primarily focused on physiological measurements of immunity or oxidative stress, but little is known about the molecular pathways by which testosterone could influence immunity and/or oxidative stress pathways. To further understand the transcriptomic consequences of experimentally elevated testosterone in the context of handicap hypotheses, we used previously published RNA-seq data from studies that measured the transcriptome of individuals treated with either a testosterone-filled or an empty (i.e., control) implant. Two studies encompassing three species of bird and three tissue types fit our selection criteria and we reanalyzed the data using weighted gene co-expression network analysis. Our results show that testosterone-treated individuals exhibited signatures of immunosuppression and we provide some evidence to suggest that the transcriptomic signature of immunosuppression is evolutionarily conserved between the three species. While our results provide no evidence to suggest testosterone mediates handicaps via pathways associated with oxidative stress, they do support the hypothesis that testosterone enforces the honesty of sexually-selected traits by influencing an individual’s immunocompetence. Overall, this study develops a framework for testing testosterone-mediated handicap hypotheses and provides guidelines for future integrative and comparative studies focused on the proximate mechanisms mediating sexually selected traits.

## 1. INTRODUCTION

There is a long-standing interest in understanding why sexually selected traits have evolved and one hypothesis suggests that mates have selected for traits that are costly to develop or bear (i.e., the handicap hypothesis; Zahavi, 1975). Under this framework, an individual’s investment in sexually selected traits is thought to correlate with their investment in traits that influence their survival (Grafen, 1990; Andersson, 1994). Individuals face tradeoffs when fitness-related traits exhibit negative correlations and, because of these tradeoffs, individuals can incur survival costs from their reproductive investments (Stearns, 1992). These costs are thought to arise because the development and/or expression of traits important for reproduction (e.g., sexually selected traits) and traits important for survival (e.g., immune function) are dependent on the same mechanism (Zera and Harshman, 2001). Therefore, our understanding of the evolution of sexually selected traits is dependent upon our understanding of the mechanisms that underlie their production and the pleiotropic effects of such mechanisms (Kokko et al., 2003).

Testosterone is a sex steroid that is known for its influence on a diverse suite of behavioral, morphological, and physiological traits associated with reproduction (e.g., metabolism and courtship behaviors; Ketterson et al. 2009), including the development and/or expression of sexually selected traits (Hau, 2007; Fusani, 2008; Ball and Balthazart, 2009). In combination with its effects on traits associated with survival (e.g., immune function, Segner et al., 2017), testosterone is thought to enforce the honesty of sexually selected traits (Ketterson and Nolan, Jr., 1999, Buchanan et al., 2001, Wingfield et al., 2001, Reed et al., 2006). Two prominent hypotheses have been proposed to explain how testosterone maintains the honesty of sexually selected traits: the Immunocompetence Handicap Hypothesis (Folstad and Karter, 1992) and the Oxidation Handicap Hypothesis (Alonso-Alvarez et al., 2007). The Immunocompetence Handicap Hypothesis proposes that sexually selected traits remain honest because of testosterone’s antagonistic effects on an individual’s immune function. Therefore, poor quality or low condition individuals cannot maintain high levels of circulating testosterone due to testosterone’s immunosuppressive effects (Folstad and Karter, 1992). An initial meta-analysis revealed weak support for this hypothesis (Roberts et al., 2004), but a subsequent meta-analysis found that experimentally increasing testosterone results in suppression of both cell-mediated and humoral immunity (Foo et al., 2017). Foo et al. (2017) also found positive trends between multiple measures of immune function and naturally occurring levels of circulating testosterone. These results fit the predictions of the Immunocompetence Handicap Hypothesis because individuals naturally expressing high of testosterone represent high quality or high condition individuals that can invest in sexually selected traits without compromising their immune system (Peters, 2000). The Oxidation Handicap Hypothesis, on the other hand, states that sexually selected traits remain honest because testosterone increases an individual’s susceptibility and/or exposure to oxidative stress (Alonso-Alvarez et al., 2007). In other words, testosterone may influence an individual’s ability to protect or repair cellular machinery from oxidative damage (e.g., an individual’s antioxidant defenses) or testosterone may influence the rate that reactive oxygen species are produced (Alonso-Alvarez et al., 2007). Importantly, either one of these consequences may occur independent of the other. Of the few studies that have directly tested the Oxidation Handicap Hypothesis, some have found support (Mougeot et al., 2009; Hoogenboom et al., 2012) while other studies did not find support for this hypothesis (Isaksson et al., 2011; Casagrande et al., 2012; Taff and Freeman-Gallant, 2014; Baldo et al., 2015). Nonetheless, both hypotheses have primarily been tested using physiological measurements of oxidative stress and immunity, but less is known about the underlying molecular pathways. Given that sex steroids partly function by binding to intracellular receptors and acting as transcription factors (Ketterson and Nolan, Jr., 1999; Nelson, 2011), measuring the relationship between testosterone and transcription can shed light on the proximate pathways that testosterone influences.

Modern sequencing approaches, like RNA sequencing (RNA-seq), allow for comprehensive measurements of whole transcriptomes and the relative abundance of each transcript (Wang et al., 2009). This approach assesses coordinated, large-scale transcriptional responses rather than focusing on targeted candidate genes (e.g., via qPCR). RNA-seq approaches have been used to investigate the role of androgens on gene expression, particularly in the context of sex differences (Gao et al., 2015; Cox et al., 2017) and gonadal development (Monson et al., 2017; Zheng et al., 2019). Similarly, RNA-seq based studies have been crucial in providing a more comprehensive understanding of the complex and dynamic immune and stress responses (e.g., Barshis et al., 2013; Huang et al., 2013; Kim et al., 2018). In the context of mate choice, measuring the relationship between testosterone and transcription can shed light on the pathways that testosterone influences to potentially enforce the honesty of sexually selected traits (e.g., immune or oxidative stress pathways). Such studies can ultimately inform our understanding of the effects of circulating testosterone on the expression of genes related to immune function and/or oxidative stress. To date, these hypotheses have rarely been tested using genome scale approaches. However, in red grouse (*Lagopus lagopus scoticus*), testosterone treatment had little effect on overall gene expression in the liver and spleen but did result in the down-regulation of genes related to immune function in caecal tissue (Wenzel et al., 2013).

Given that Wenzel et al. (2013) used a microarray-based approach, testing these handicap hypotheses using RNA-seq represents a more modern, robust test as RNA-seq provides many advantages over microarray technologies, including higher sensitivity and no hybridization biases (Wang et al., 2009). Here, we use published RNA-seq datasets to further examine the effects of testosterone on the transcriptome. We re-analyze studies that measured the transcriptome of testosterone-treatment and control subjects in three species of bird: zebra finch (*Taenopygia guttata*), golden-collared manakin (*Manacus vitellinus*), and Japanese quail (*Coturnix japonica*). In the zebra finch, male vocal behavior as well as bill color (a sexually-selected trait) have been found to be sensitive to testosterone (Cynx et al., 2005, McGraw et al. 2006). Similarly, male golden-collared manakin engage in elaborate courtship behaviors during the breeding season and experimentally blocking androgen receptors decreases the expression of male courtship behaviors (Day et al., 2007; Schlinger et al., 2013). Male Japanese quail also produce brightly colored cheek feathers to attract females (Hiyama et al., 2018). Castrating males influences the color of a male’s cheek feathers and administering testosterone to castrated males causes cheek patches of castrated males to match those of males that have not been castrated. In this study, we focus on transcriptomic data from tissues that are known to be express androgen receptors: the foam gland of quail and muscular tissues from the zebra finch and golden-collared manakin, (Adkins-Regan, 1999; Fuxjager et al., 2016). We constructed co-expression networks to identify gene networks that show correlated expression patterns following testosterone treatment. If testosterone influences the honesty of sexually selected traits via immune mechanisms, relative to the control group, the group treated with testosterone treatment will exhibit lower expression of genes with annotated immune function (i.e. immunosuppression). Alternatively, if testosterone acts via mechanisms associated with an individual’s susceptibility or exposure to oxidative stress, then we predict that testosterone treatment will cause a decrease in the expression of genes with annotated functions in antioxidant protection and/or an increase in genes that are expressed in response to oxidative stress. An important caveat is that no support for either hypothesis does not exclude the possibility that these pathways maintain the honestly of sexually selected traits independent of testosterone’s effects, as has been suggested before (Metcalfe and Alonso-Alvarez 2010, Weaver et al. 2017). Lastly, given the limited taxonomic diversity of our dataset, we acknowledge that future studies including more diverse taxon and tissues are ultimately needed to robustly examine if testosterone mediates the honesty of sexually selected traits via transcriptomic pathways.

## 2. METHODS

### 2.1. Study Selection

To identify studies of interest, we first performed a literature search on both Scopus and Google Scholar with the following search terms: “testosterone” AND “RNA-seq” or “transcriptome” or “transcriptomics”. This literature search produced 260 results. From this list of 260 studies, we retained RNA-seq studies that measured gene expression in adult males from both testosterone-manipulated and control groups. This process resulted in one study for re-analysis and we also identified an additional dataset by searching within NCBI’s Sequence Read Archive (Supplemental Figure 1). Finseth and Harrison (2018) experimentally increased testosterone in Japanese quail (*Coturnix japonica*, “quail”) experiencing short days and performed RNA-seq on the foam gland (n=6 each testosterone and control). Fuxjager et al. (2016) experimentally increased testosterone in male golden-collared manakins (*Manacus vitellinus*, “manakin”) and performed RNA-seq on pectoralis and scapulohumeralis caudalis tissue (n = 3 individuals per treatment group and 2 tissue types per individual). In an effort to understand the role of androgen receptors in mediating the gene expression patterns observed in the manakin samples, Fuxjager et al. (2016) also administered 2 different combinations of hormone implants to adult zebra finches (*Taeniopygia guttata*): one group received testosterone-filled implants and an empty implant while the second group was administered a testosterone implant and an implant filled with flutamide, a drug that blocks the androgen receptor. Fuxjager et al. (2016) then measured the transcriptome of the pectoralis and scapulohumeralis caudalis muscles (n = 3 individuals per treatment group and 2 tissues types per individual) using RNA-seq. Thus, no unmanipulated, control zebra finch samples were included in the zebra finch RNA-seq data generated by Fuxjager et al. (2016). Because the current study is not focused on understanding the role of the androgen receptor in mediating handicap hypotheses and due to the limited sample size available for testing the role of androgen receptor in mediating handicap hypotheses, only samples from the zebra finches treated with just testosterone were included in our analysis. While the zebra finch samples lack an appropriate control (i.e., individuals not treated with testosterone), the zebra finch samples can nonetheless be used in the shared orthologous gene expression analysis (see below for more details).

**Figure 1.**
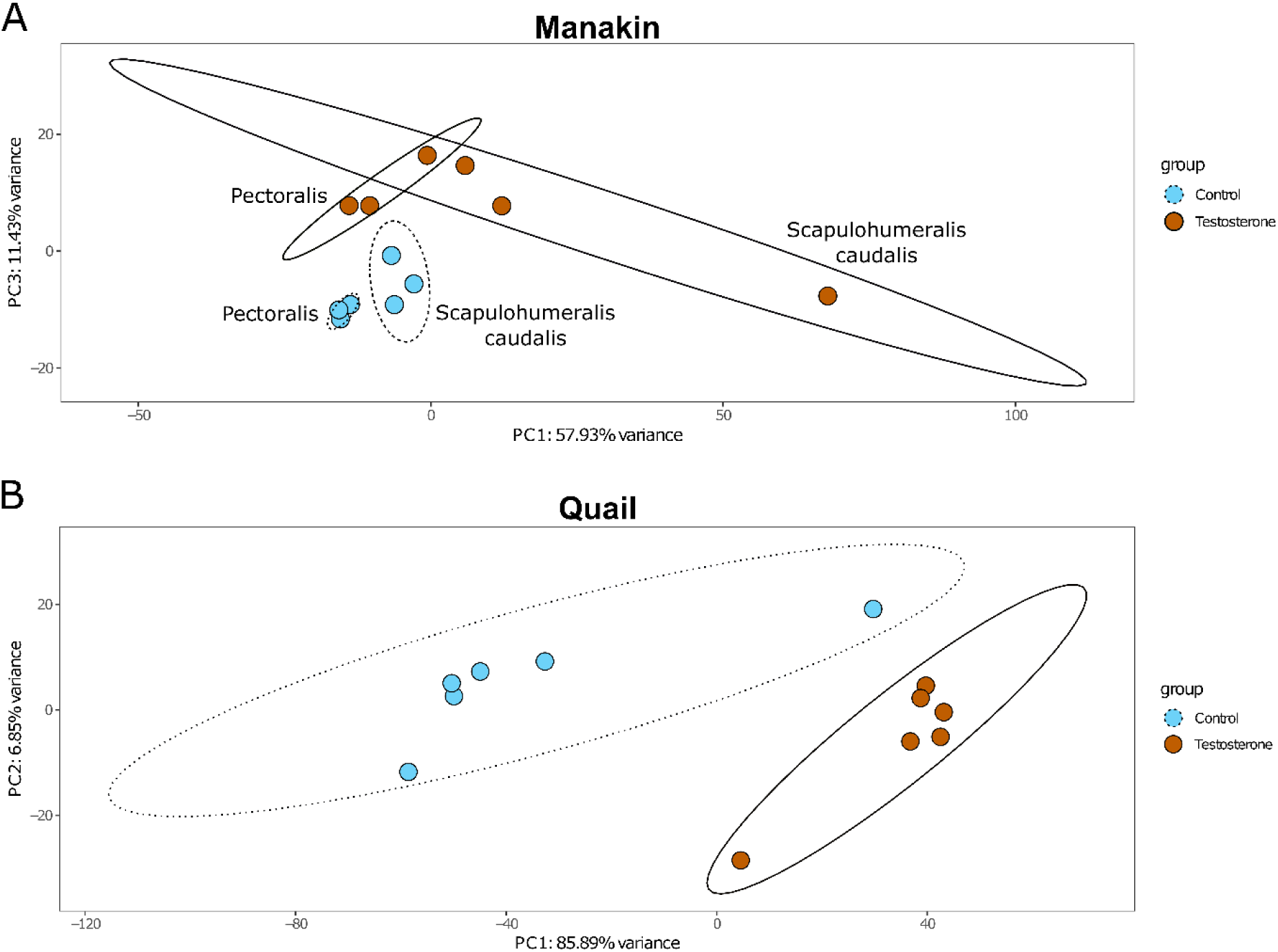
PCA of (A) manakin and (B) quail. Samples separate by treatment along PC3 for manakin and PC1 for quail. Each circle represents a sample and is color-coded by treatment. Manakin samples are labeled by muscle type. Ellipses represent 95% confidence intervals.

### 2.2 Data Re-analysis

We downloaded the raw sequencing data from SRA with sratoolkit fastq-dump (quail: PRJNA397592; manakin & zebra finch: PRJNA297576) and adaptor trimmed all reads with Trim Galore! v0.3.8 (https://github.com/FelixKrueger/TrimGalore). We aligned trimmed reads to the respective reference genome (*M. vitellinus* v2, *C. japonica* v2, *T. guttata* bTaeGut1_v1.p) for each species with STAR v2.5.3 (Dobin et al., 2013) and quantified expression with htseq-count v0.6.0 (Anders et al., 2015), specifying strand ‘no’. We normalized counts to sequencing depth and variance stabilizing transformed counts with DEseq2 (Love et al., 2014). Transformed counts were visualized with a principal component analysis (PCA) using pcaExplorer v2.8.1 (Marini & Binder 2019) and PCAtools v2.0.0 (Blighe & Lun 2020).

To test for the effect of testosterone treatment on transcription, we used the weighted gene co-expression network analysis (WGCNA) tool (Langfelder et al., 2011; Langfelder and Horvath, 2008). We created modules independently for quail and manakin with the following shared parameters: network type=signed, minimum module size=30, and module dissimilarity=0.2. We used β=12 for quail and β=18 for manakin, which represents the point the network reached scale free topology. We then tested for correlations between modules and testosterone treatment using a p < 0.05 cutoff. We identified the hub genes of each module by selecting the top five genes with the highest module membership (MM) score. Since we created a signed network in WGCNA and correlated modules to a binary trait of Testosterone or Control, each module correlated with testosterone treatment shows opposite expression patterns between the two treatment groups. In other words, genes showing low expression in testosterone treated animals are relatively highly expressed among control samples.

As each of these studies was relatively underpowered for gene expression analyses (reviewed in Schurch et al. 2016), we used a subsampling approach to test the robustness of each WGCNA analysis. In other words, we were interested if the results obtained in the full analysis were reproducible using subsets of the data. Gene signatures observed in both the full analysis and subset analysis would thus help confidently identify meaningful pathways responding to testosterone treatment. While both studies contain the same number of RNA-seq samples, the manakin dataset represents 6 total individuals and the quail dataset represents 12 individuals. We divided the quail data into two sets of six individuals (n=3 control, n=3 testosterone). For manakin, we divided the data in two by muscle type (n=3 control, n=3 testosterone each). We then re-ran WGCNA on each of the four subsets with the same parameters, except specifying β=16 for both quail subsets and β=18 for both manakin subsets. We then performed functional enrichment analyses on each module as described below.

We were also interested if there was a shared response to testosterone among all species. Therefore, using all 30 RNA-seq samples (n=12 quail, n=12 manakin, n=6 zebra finch), we identified 8,454 one-to-one (i.e. single copy, non-duplicated) orthologs among all samples using Ensembl BioMart (Kinsella et al. 2011). This filtering process restricts the analyses to genes that are evolutionarily conserved between species. We normalized and variance stabilized transformed all samples using DESeq2. Since the data come from two independent studies, we then used the removeBatchEffect function from limma (Ritchie et al., 2015) to control for batch effects. Lastly, we used these conserved, expressed orthologs to create a single co-expression network with WGCNA for all samples. Module reconstruction parameters were identical to the species-specific analysis described above, except we used β=12.

To identify functional pathways in each of the WGCNA modules, we performed an Overrepresentation Analysis with WebGestaltR (Liao et al., 2019) using the Gene Ontology Biological Processes database. For each module, we tested the genes found in each module against a background list of all genes used to perform the WGCNA analysis. GO categories were significantly enriched if the FDR value < 0.05. We then interrogated significant GO categories, with an emphasis on immune and oxidative stress related pathways, in modules significantly correlated with testosterone treatment. To find support for the Immunocompetence Handicap Hypothesis, immune related categories (e.g., “immune system process”) had to be significantly enriched among genes expressed lower in testosterone treated animals. To find support for the Oxidation Handicap Hypothesis, oxidative stress related GO categories had to be significantly enriched among genes highly expressed (e.g., “response to oxidative stress”) or gene exhibiting lower expression levels (e.g., “antioxidant activity”) following testosterone treatment.

## 3. RESULTS

After filtering, we used 13,509 manakin genes and 13,946 quail genes for PCA and WGCNA network construction. Testosterone treatment had pronounced effects on gene expression and individuals clustered by treatment in both species-specific comparisons (Figure 1) and in the analysis ran on orthologous genes (Supplemental Figure 2).

**Figure 2.**
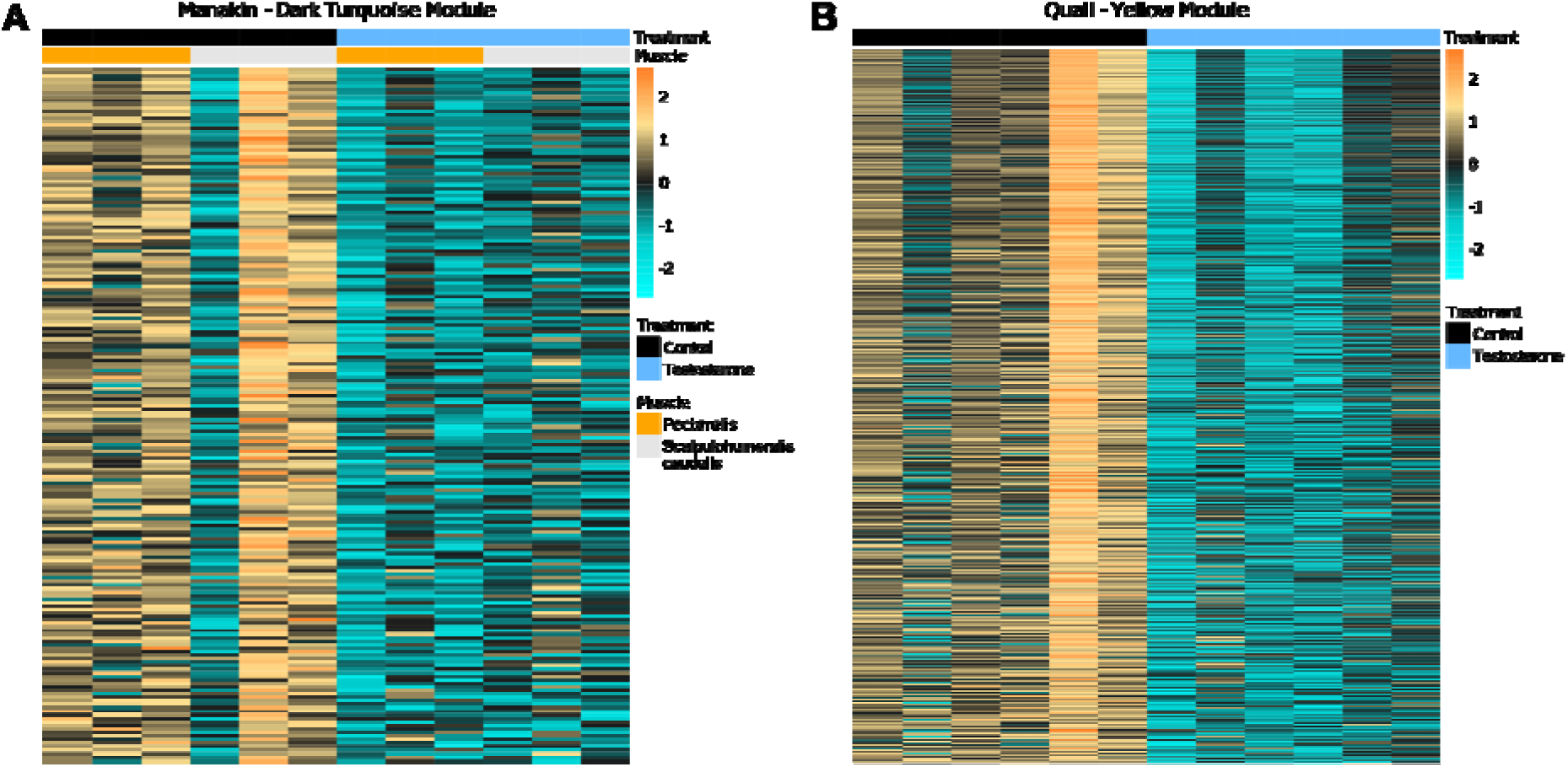
Expression heatmaps of the (A) Manakin Dark Turquoise Module and (B) Quail Yellow Module, which represent low expression of immune related genes in testosterone, but not control, treated animals. Each column represents a sample color coded by treatment or muscle type. Each row represents a module gene. High expression is indicated by orange colors and low expression is represented by blue colors.

### 3.1 WGCNA – Quail

WGCNA constructed 18 modules for quail, six of which were correlated with testosterone treatment (Supplemental Figure 3). The yellow module (925 genes, r=-0.74) was strongly enriched for broad immune related GO categories spanning both the innate (e.g. “inflammatory response”) and adaptive (e.g. “T cell activation”) arms of the immune system (Table 1, Supplemental Table 1). This represents a significant decrease in immune gene expression following treatment (Figure 2A). The yellow module hubs were *SASH3, ITGB2, SLAMF8* (LOC107324444), *TRAF3IP3*, and *EVI2A*.

**Table 1.** GO enrichment for modules (from both species) that are associated with immune function. The top 5 gene ontology (GO) categories are presented, along with FDR adjusted p-value and WebGestaltR enrichment score.

| GO ID | Description | FDR | Enrichment |
| --- | --- | --- | --- |
| *Quail, Yellow Module* |  |  |  |
| GO:0050778 | positive regulation of immune response | 0 | 3.52 |
| GO:0001816 | cytokine production | 0 | 3.36 |
| GO:0046649 | lymphocyte activation | 0 | 4.52 |
| GO:0045087 | innate immune response | 0 | 3.02 |
| GO:0019221 | cytokine-mediated signaling pathway | 0 | 3.47 |
| *Manakin, Dark Turquoise Module* |  |  |  |
| GO:0030217 | T cell differentiation | $6.56 \times 10^{-12}$ | 12.78 |
| GO:0002521 | leukocyte differentiation | $6.56 \times 10^{-12}$ | 7.83 |
| GO:0030098 | lymphocyte differentiation | $2.27 \times 10^{-11}$ | 9.69 |
| GO:0042110 | T cell activation | $1.57 \times 10^{-10}$ | 8.00 |
| GO:0030097 | hemopoiesis | $6.11 \times 10^{-9}$ | 4.99 |

**Figure 3.**
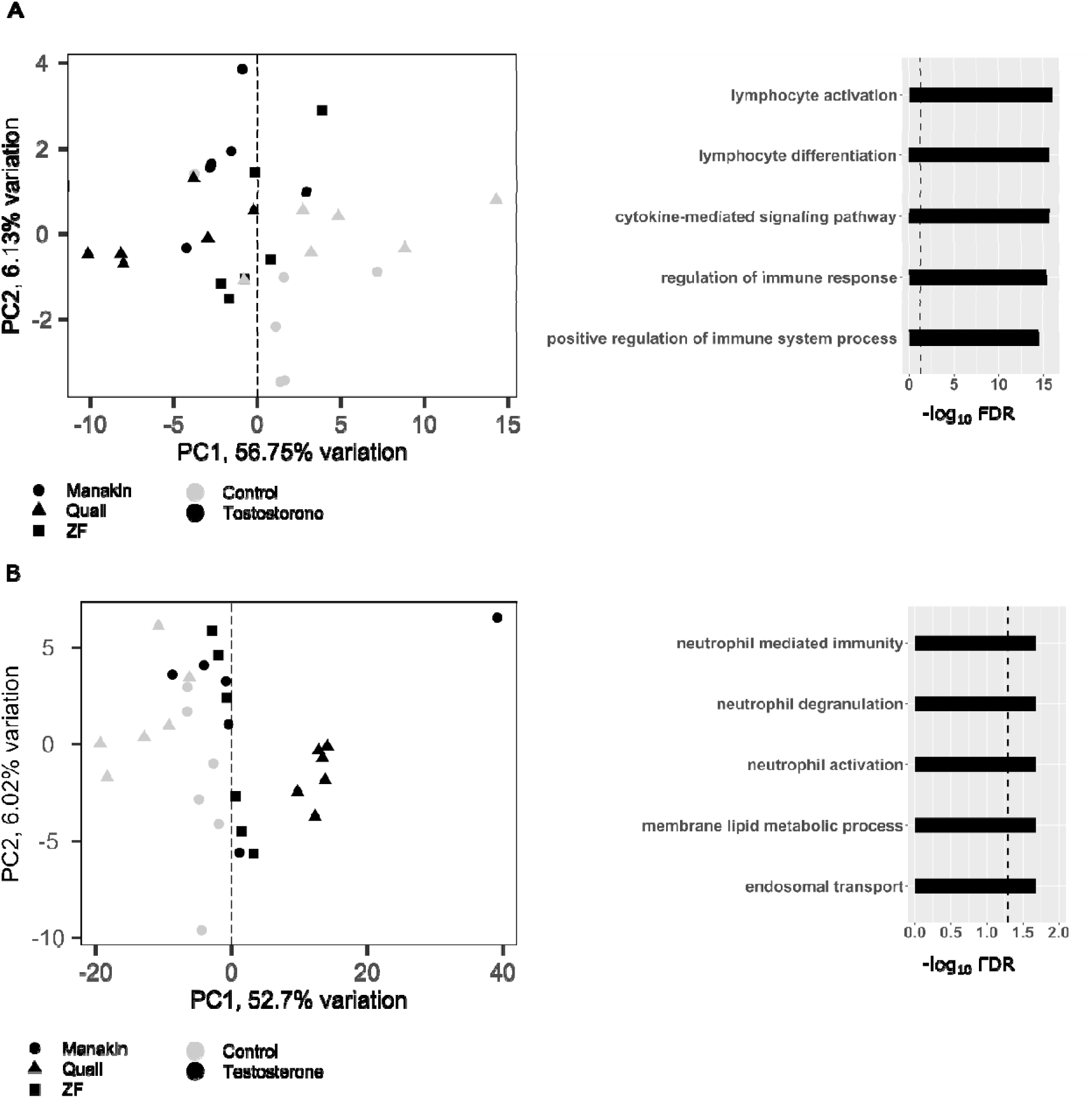
PCA and functional enrichment of the (A) purple and (B) black modules from the orthologous gene WGCNA. PCAs were constructed using genes belonging to each module. The shape of the points represents species and the color represents treatment. The dashed line in the enrichment bar plots represents FDR=0.05.

The purple and black modules were also negatively correlated with testosterone treatment and were enriched for translation and muscle process GO categories respectively. Additionally, we found two modules containing genes highly expressed following testosterone treatment. The turquoise module was the most strongly correlated with testosterone treatment (4423 genes, r=0.98). GO enrichment was largely driven by genes involved in the Golgi apparatus and endoplasmic reticulum functions (Supplemental Table 1). The green module (795 genes, r=0.61) was primarily enriched for broad transcriptional regulation and protein modification processes. Lastly, we found no modules that were associated with either antioxidant defenses or oxidative damage.

### 3.2 WGCNA – Manakin

WGCNA constructed 34 modules for manakin, 12 of which were correlated with testosterone treatment (Supplemental Figure 3). Seven modules were correlated with muscle type but, given none of these seven modules were also correlated with testosterone treatment, our results provide no support for a tissue specific response to testosterone at the network level. Of the 12 modules, 7 were negatively correlated and 5 positively correlated with testosterone treatment. Like the quail, manakins treated with testosterone also exhibited a significant decrease in immune gene expression relative to control individuals (Figure 2B, Supplemental Table 2). The dark turquoise module (198 genes, r=-0.71) was strongly enriched for a broad range of immune related GO categories spanning both the innate and adaptive components of the immune system (Table 1, Supplemental Table 2). The dark turquoise hub genes were *MHC1A* (LOC108639055), *INPPL1* (LOC103767762), *CCL14* (LOC103758017), *CCL3L* (LOC103757995), and an uncharacterized non-coding RNA (LOC108640668).

The remaining negatively correlated modules were primarily enriched for glycolysis and mitochondria related categories (steel blue, pale turquoise). Among the positively correlated modules, we only observed significant GO enrichment in the light cyan module, which encompassed transcriptional regulation (Supplemental Table 2). Lastly, like the quail, we found no modules with significant enrichment of pathways related to oxidative damage or protection.

### 3.3 WGCNA – Subset Analysis

Our subsampling approach resulted in 43 modules for quail subset 1, 56 modules for quail subset 2, 55 modules for manakin subset 1, and 53 modules for manakin subset 2 (Supplemental Figure 4). Four modules in quail subset 1, six modules in quail subset 2, and three each in the manakin subsets were correlated with testosterone treatment and, of these modules, a single immune-related module was correlated with testosterone-treatment in both quail subsets and one manakin subset. Both the greenyellow module (n=518 genes) in quail subset 1 and antiquewhite1 module (n=1280 genes) in quail subset 2 were primarily enriched for lymphocyte activation and T cell signaling pathways. Two hundred & fifty-two genes were shared between these two quail modules and these genes were enriched for broad immune system processes, including lymphocyte and T cell activation. Combined, the greenyellow and antiquewhite1 modules contain 688/925 genes found in the quail yellow module from the full analysis. The blanchedalmond module (n=102 genes) in manakin subset 1, which represents pectoralis tissue, was primarily enriched for interleukin and cytokine signaling (Supplemental Table 3). This module contained 41/198 genes found in the dark turquoise module from the full analysis of manakin data. In each case, the module consisted of immune related genes exhibiting lower expression levels in the testosterone treated group relative to the control group (Supplementary Table 3). Manakin subset 2, which represents scalpulohumeralis caudalis tissue, did not have an immunosuppression signature. None of the subsets exhibited an oxidative stress signature correlated with testosterone treatment.

### 3.4 WGCNA – Shared response among Quail, Manakin, & Zebra Finch

Our integration of all three species resulted in 18 modules, 8 of which were correlated with testosterone treatment in all three species (i.e., p < 0.05; Supplemental Figure 5; Supplemental Table 4). Of these testosterone responsive modules, the purple module (195 genes, r = -0.58) was significantly enriched for broad immune related categories (Figure 3A). This result represents lower expression of these module genes in testosterone treated animals (relative to control animals) and reflects the shared response of immunosuppression among all three species. Interestingly, the black module (1,331 genes, r = 0.58) also showed significant enrichment of neutrophil mediated immunity, which represents increased expression of these module genes in testosterone treated animals (Figure 3B). This analysis reveals that while testosterone leads to widespread immunosuppression, a small component of the immune system is also activated.

## 4. DISCUSSION

In this study, we examined transcriptomic responses to experimentally increased circulating testosterone in three species of bird. As expected based on the pleiotropic nature of sex steroids (Ketterson and Nolan, Jr., 1999, Hau 2007), our gene network analyses demonstrate that, in all three species, individuals treated with testosterone exhibited altered expression patterns of a diverse suite of genes relative to controls (e.g., regulation of immune response, hemopoiesis, endosomal transport). In the context of handicap hypotheses, the expression genes associated with immune function was significantly lower in the testosterone-treated group lower relative to controls, providing transcriptomic support for the Immunocompetence Handicap Hypothesis.

These results also begin to elucidate the molecular pathways (e.g., lymphocyte activation, cytokine signaling, T cell activation) potentially underlying the testosterone-induced immunosuppression previously documented using physiological measures of immune function in Foo et al. (2017). Moreover, testosterone is necessary to produce the secondary sexual characteristics associated with mating in all three species (Cynx et al., 2005, McGraw et al. 2006, Schlinger et al., 2013, Hiyama et al., 2018). Our results therefore begin to detail the potential molecular pathways underlying the trade-off between the expression of sexually selected traits and immune function in these species. We did not find support for the Oxidation Handicap Hypothesis as there was no enrichment of genes expressed related to oxidative damage, nor suppression of genes related to antioxidant defenses in either species. These results suggest that costs associated with maintaining high levels of circulating testosterone are not via pathways associated with an individual’s susceptibility or exposure to oxidative stress. Importantly, oxidative stress could still be involved in enforcing the costs of reproduction or sexually selected traits, but our results suggest that this cost is not borne out via molecular pathways that are sensitive to testosterone, at least in the limited number of tissues and species examined here.

Our analyses revealed that transcriptomic immunosuppression was broad, encompassing aspects of both innate immunity (e.g., interferon signaling and cytokine signaling) as well as adaptive immunity (e.g., antigen processing and presentation; Table 1, Supplemental Tables 1 & 2). While the observed effect of testosterone could occur through non-genomic pathways, previous work examining the relationship between androgen receptors and the immune system suggests genomic pathways are involved (Trigunaite et al., 2015; Segner et al., 2017; Gubbels Bupp and Jorgensen, 2018). More specifically, while testosterone exposure and subsequent androgen receptor activity can promote innate immune cell differentiation and development, testosterone also reduces activity of these cells (Gubbels Bupp and Jorgensen, 2018). Indeed, the hub genes of the immune related modules highlight broad suppression of innate immune signaling (quail yellow: *SASH3, SLAMF8, TRAF3IP3*; manakin dark turquoise: *INPPL1, CCL14, CCL3L*, ncRNA; (Beer et al., 2005; Veillette, 2010; Dauphinee et al., 2013; Sokol and Luster, 2015; Zou et al., 2015; Thomas et al., 2017; Wang et al., 2018). Similarly, testosterone impacted expression of genes involved in the adaptive immune system. Testosterone exposure reduces T cell activity, a crucial component of proper adaptive immune signaling, (Lin et al., 2010; Kissick et al., 2014), and this is a prominent signature in both quail (Supplemental Table 1) and manakin (Table 1). In addition to suppression of T cell activity in manakin, we also identified MHC class IA as a hub gene in the manakin dark turquoise module. MHC class IA binds and presents viral peptides to CD8+ T cells, which is a critical component of the adaptive immune response (Neefjes et al., 2011). Previous work has shown suppressive effects of testosterone on CD4+ T cells/MHC class IIB (Lin et al., 2010) and CD8+ T cells (Page et al., 2006). However, our study is the first to describe suppression of genes involved in T cell activity as well as MHC class I.

The subset analysis demonstrated that small sample sizes can influence both module construction and functional enrichment. For quail, we identified a single module in each subset that was significantly correlated with testosterone treatment and enriched for immune pathways. As with the original full quail analysis, both immune related modules from the quail subsets exhibited enrichment for T cell related activity, suggesting that testosterone has a robust effect on T cell signaling in the quail foam gland (Supplemental Table 3). In contrast, we identified a single immune related module correlated with testosterone treatment only in the manakin subset containing pectoralis tissue. Whereas the full manakin analysis revealed a T cell signaling signature, the subset analysis primarily exhibited a cytokine production and signaling signature. Individual variation in gene expression, tissue-specific expression, responses to the treatment, as well as the limited sample size likely influenced our ability to detect an immunosuppression signature in both manakin subsets. Indeed, the manakin pectoralis subset exhibited a cytokine signature not observed in the full analysis, suggesting individuals and/or muscle types exhibited quite variable responses to testosterone treatment. Overall, the results of the subset analysis highlight the sensitivity of WGCNA to small sample sizes and underscore the importance of adequate sample sizes for future studies using this analytical approach.

As with each species-specific analysis, the WGCNA conducted on orthologous genes (i.e., evolutionary conserved genes) also identified a broad signature of immunosuppression in the testosterone-treated group but not the control groups. This result provides preliminary insight into the evolutionary conserved transcriptomic response to testosterone and provides additional support for the immunocompetence handicap hypothesis. The increased power of the WGCNA conducted on the orthologous genes (n=30 samples) may have reduced the influence of pre-existing individual differences and allowed us to detect more subtle changes in gene expression that were missed in the species-specific analyses (n=12 samples). For instance, a module encompassing genes related to neutrophil-mediated immunity was highly expressed following testosterone treatment. Neutrophil activity, a hallmark of innate immunity, is associated with inflammation predominantly in lymphoid tissues, but their broader tissue-specific functions remain uncertain (reviewed in Ng et al., 2019). Interestingly, we also observed an inflammation signature within the purple module (Supplemental Table 4), which contains genes exhibiting lower expression levels in the testosterone treatment relative to controls. Overall, these results suggest the changes in inflammation-related gene expression following testosterone treatment are dynamic and that studies with larger sample size are better able to detect more complex responses.

Our results were based on the transcriptomes from skeletal muscles or the quail foam gland, tissues which are not traditionally studied in avian immunology (Rose, 1979; Schat et al., 2014). A growing body of research on fish and mammals does however suggest that muscle tissues play an important role in immune responses (reviewed in Valenzuela et al. 2017).

Nonetheless, in combination with the testosterone-related patterns of immunosuppression found in Foo et al. (2017), our study provides some additional evidence to suggest that RNA-seq can detect functional signatures in non-traditional tissues (e.g., Louder et al., 2018). As such, public repositories of genomic data will become increasingly valuable to test evolutionary hypotheses in novel ways (e.g. Shultz & Sackton 2019).

## 5. CONCLUSION AND FUTURE DIRECTIONS

Testosterone-mediated handicap hypotheses have captivated biologists since their development but have primarily been tested using physiological measures of oxidative stress or immune function (e.g., Roberts et al. 2004, Foo et al. 2017). Here we develop a transcriptomic approach for testing these hypotheses. Our findings, while limited by sample size and taxonomic diversity, document species-specific and evolutionary conserved signatures of immunosuppression in three species of bird. These results provide a foundation for future transcriptomic studies encompassing additional tissue types and/or more taxonomically diverse species. Future studies should prioritize measuring testosterone’s effect on gene expression using a within-individual sampling approach to mitigate the influence of pre-existing individual differences in gene expression. Studies should also strive for sample sizes > 20, as recommend by the developers of WGCNA (Langfelder & Horvath 2017). The importance of larger sample sizes is exemplified here by the subset analysis which shows that modules built using smaller sample sizes recapitulate an immunosuppression signature most but not all the times (i.e., 3 out of 4 subsets included immune-related modules significantly correlated with treatment in this study).

Additionally, the immune modules within the 3 subsets exhibited variable immunological pathway enrichment compared to the full analysis potentially due to individual variation influencing the genes found in each module. When terminal sampling is the only option for collecting samples, a common scenario for transcriptomic studies, larger sample sizes are needed to reduce the influence of such pre-existing individual differences. Furthermore, future works should also consider conducting experimental infections and/or immune challenges in combination with RNA-seq analyses to examine how transcriptomic signatures relate to immune function. Novel endocrine-based experiments similar to Goymann et al. (2015) and Goymann and Dávila (2017) paired with RNA-seq analyses can also shed light on the transcriptomic responses to acute stimulation of the hypothalamic-pituitary-gonadal axis (i.e., the endocrine cascade associated with gonadal testosterone production). Future transcriptomic studies should also consider the best practices for experimental design to ensure appropriate sample sizes are selected and that any results are biologically meaningful (Schurch et al. 2016). Overall, these integrative, comparative, and mechanistic approaches will ultimately provide novel insights into the mechanisms underlying handicap hypotheses and, more broadly, the evolution of sexually selected traits.

## ACKNOWLEDGEMENTS

We would like to thank Ignacio Moore, Dana Hawley, Heather Watts, Christopher Balakrishnan, and three anonymous reviewers for providing helpful feedback on earlier versions of this manuscript.

