## Supplemental Figures for "Developing a transcriptomic framework for testing testosterone-mediated handicap hypotheses"

**Supplemental Figure 1. Workflow to identify studies of interest for reanalysis. Numbers next to arrows represent the number of studies retained at each step.**

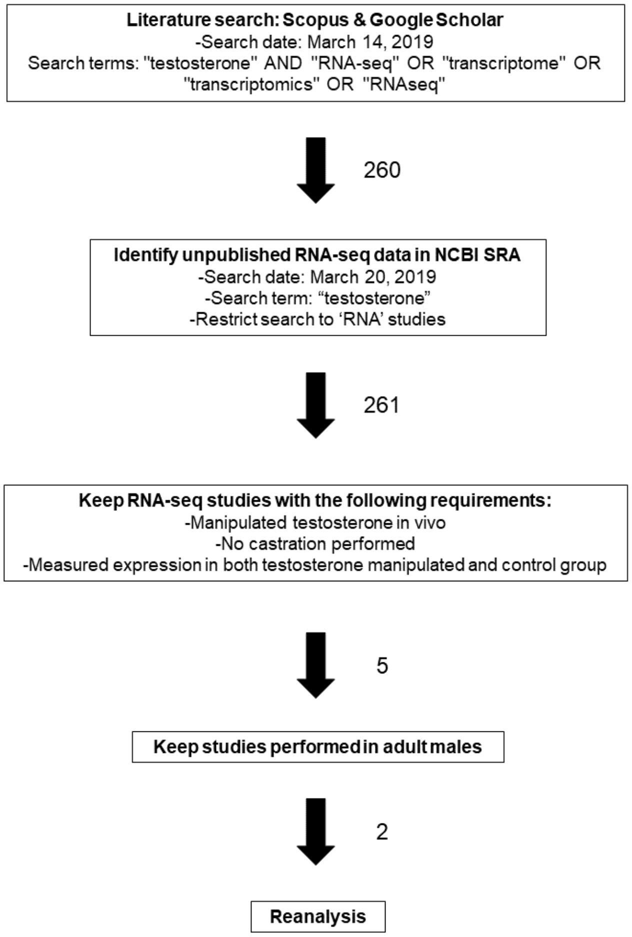

**Supplemental Figure 2. PCA of all 8454 orthologous genes. Samples are colored by treatment and shaped by species.**

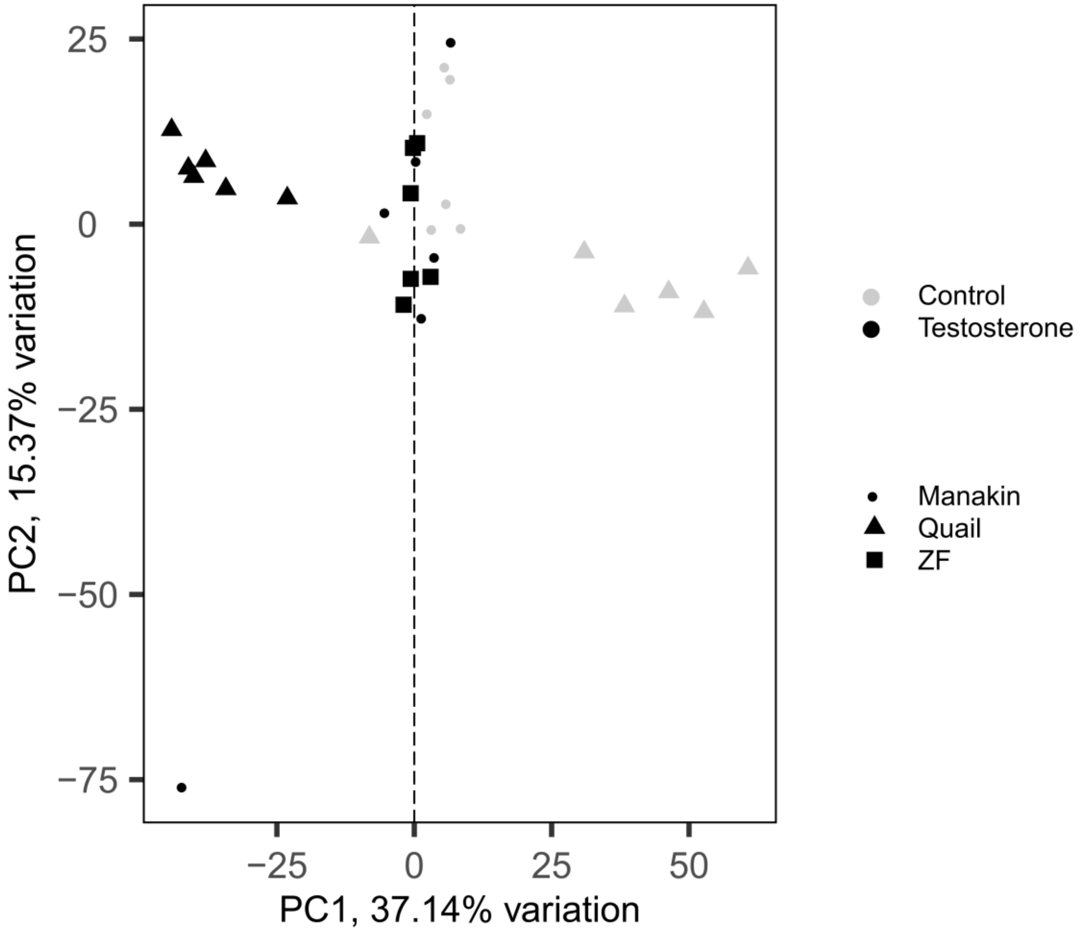

**Supplemental Figure 3. Module trait relationships for the quail and manakin datasets.**

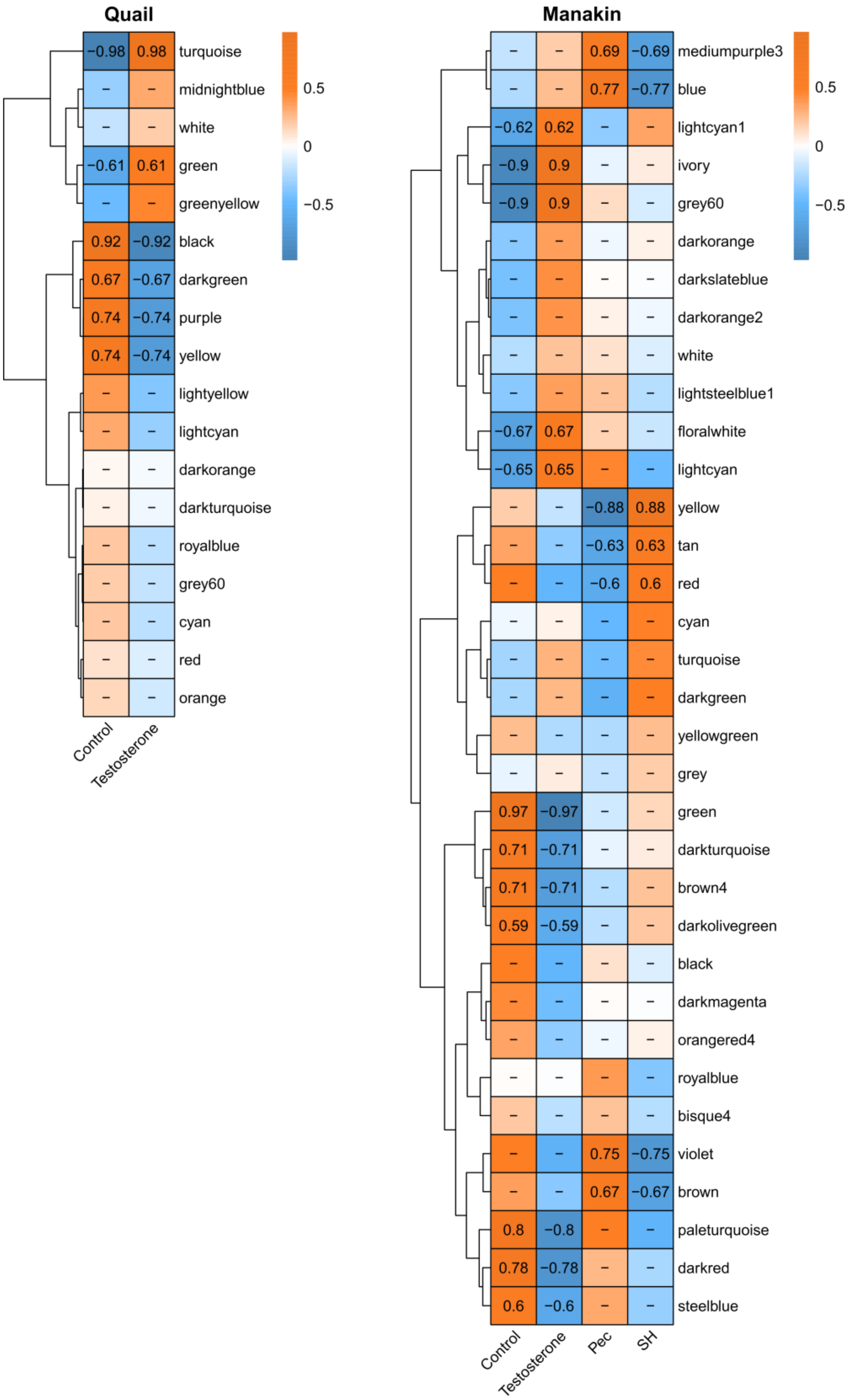

**Supplemental Figure 4. Module trait relationships for the subsampling WGCNA analysis.**

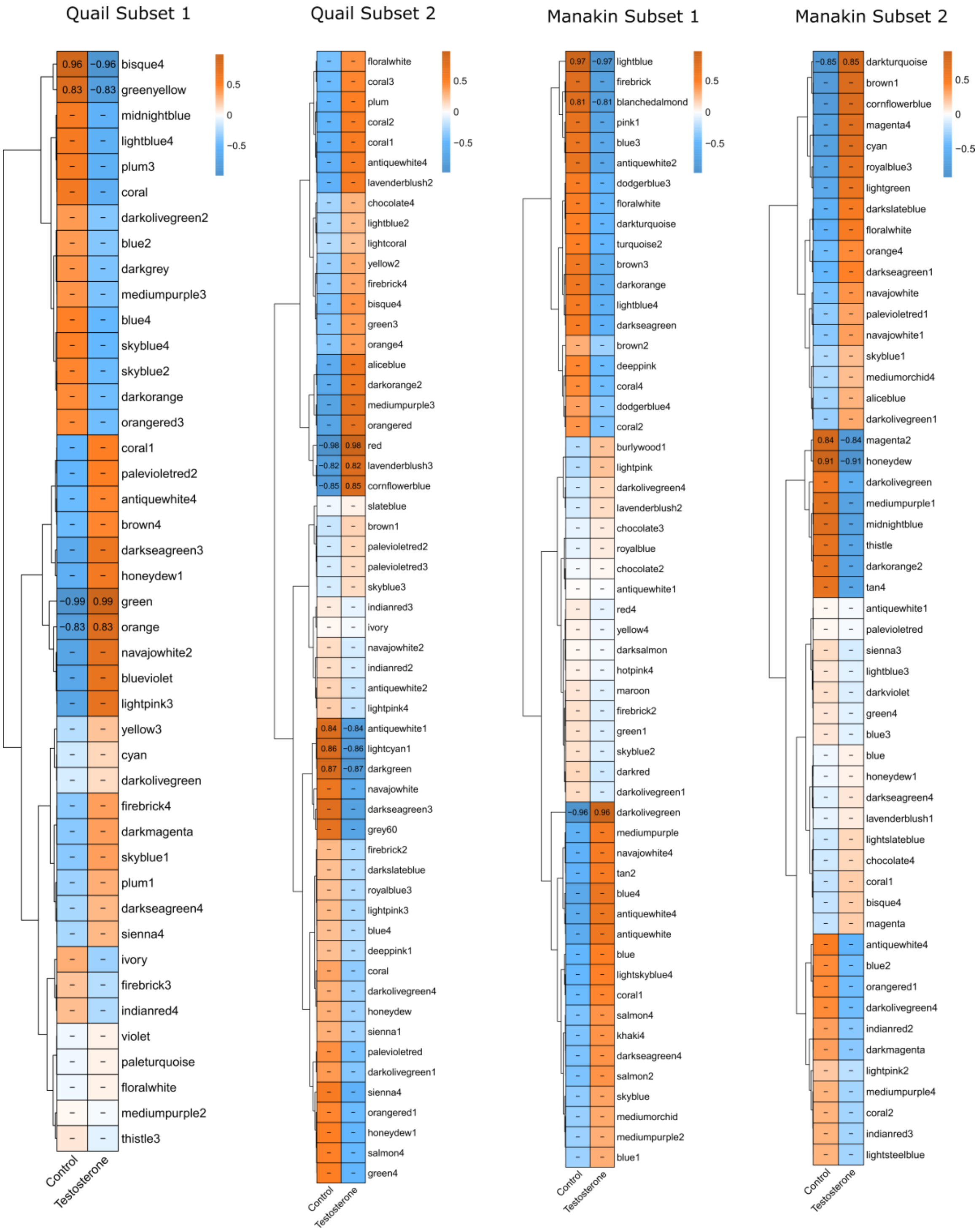

**Supplemental Figure 5. Module trait relationships for the orthologous WGCNA analysis.**

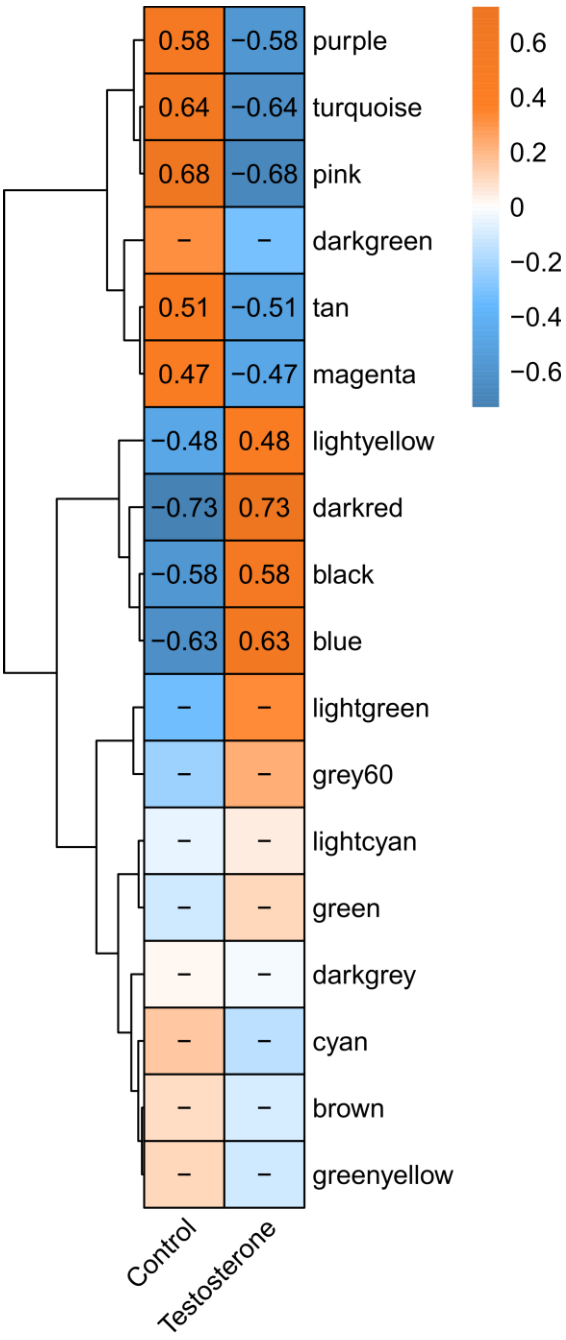
